# Identification of the *C. sordellii* lethal toxin receptor elucidates principles of receptor specificity in clostridial toxins

**DOI:** 10.1101/871707

**Authors:** Hunsang Lee, Greg L. Beilhartz, Iga Kucharska, Swetha Raman, Hong Cui, Mandy Hiu Yi Lam, John L. Rubinstein, Daniel Schramek, Jean-Philippe Julien, Roman A. Melnyk, Mikko Taipale

**Author notes:** co-first authors (Hunsang Lee, Greg L. Beilhartz, Iga Kucharska).

## Abstract

*Clostridium sordellii* lethal toxin (TcsL) is responsible for an almost invariably lethal toxic shock syndrome associated with gynecological *C. sordellii* infections. Here, using CRISPR/Cas9 screening, we identify semaphorins SEMA6A and SEMA6B as the cellular receptors for TcsL and demonstrate that soluble extracellular SEMA6A can protect mice from TcsL-induced edema. A 3.3 Å cryo-EM structure shows that TcsL binds SEMA6A with the same region that the highly related *C. difficile* TcdB toxin uses to bind structurally unrelated Frizzled receptors. Remarkably, reciprocal mutations in this evolutionarily divergent surface are sufficient to switch receptor specificity between the toxins. Our findings establish semaphorins as physiologically relevant receptors for TcsL, and reveal the molecular basis for the difference in tissue targeting and disease pathogenesis between highly related toxins.

---

*Clostridium sordellii* is an anaerobic gram-positive bacterium found in soil and in the gastrointestinal and vaginal tracts of animals and humans. *C. sordellii* is present in the rectal or vaginal tract of 3-4% of women, but vaginal colonization rate after childbirth or abortion is as high as 29% (*1, 2*). While the majority of carriers are asymptomatic, pathogenic *C. sordellii* infections arise rapidly and are highly lethal. Most *C. sordellii* infections occur in women following childbirth, medically induced abortion, or miscarriage, leading to a toxic shock syndrome with almost 100% mortality within days (*1–4*). The primary cause of the high mortality associated with *C. sordellii* infections is the lethal toxin TcsL (*5*), which belongs to the large clostridial toxin (LCT) family together with toxins from related species such as *Clostridium difficile* (*6*). LCTs enter the host cell by receptor-mediated endocytosis followed by translocation into the cytoplasm, where they potently modulate host cell function by glucosylating small Ras family GTPases (*6*). Although all LCTs are highly similar at the sequence level, they differ in their tissue specificity and in their effects on cell morphology, physiology, and viability. TcsL is most closely related to the *C. difficile* cytotoxin TcdB, sharing almost 90% sequence similarity. TcdB binds Frizzled family receptors FZD1/FZD2/FZD7 expressed in the colonic epithelium (*7, 8*), the primary site of *C. difficile* infection. *C. sordellii*, however, does not infect or damage colonic epithelium, suggesting that TcsL binds a different cell-surface receptor.

To identify potential cell-surface receptors for TcsL, we conducted a genome-wide CRISPR/Cas9 screen in Hap1 cells. Cells were infected with the genome-wide TKOv3 gRNA library (*9*) and treated with 0.1 nM or 1 nM TcsL. Sequencing of gRNAs from the surviving population revealed only two genes that were significantly enriched in both screens (**Fig. 1A**, fig. S1A and table S1). One was UGP2 (UDP-glucose pyrophosphorylase 2), which is required for the synthesis of UDP-glucose, the sugar donor for the glucosylation activity of all LCTs (*6*). The other was SEMA6A, encoding a transmembrane axon guidance molecule not previously linked to toxin function. Notably, despite high sequence similarity between TcsL and TcdB, our screen did not identify Frizzled receptors, or CSPG4 or PVRL3, two other TcdB-associated receptors (*7,10,11*). Neither did we identify known receptors for *C. perfringens* TpeL or *C. difficile* TcdA, other related LCTs (*12*, *13*), although all these receptors are expressed in Hap1 cells (fig. S1B).

**Figure 1.**
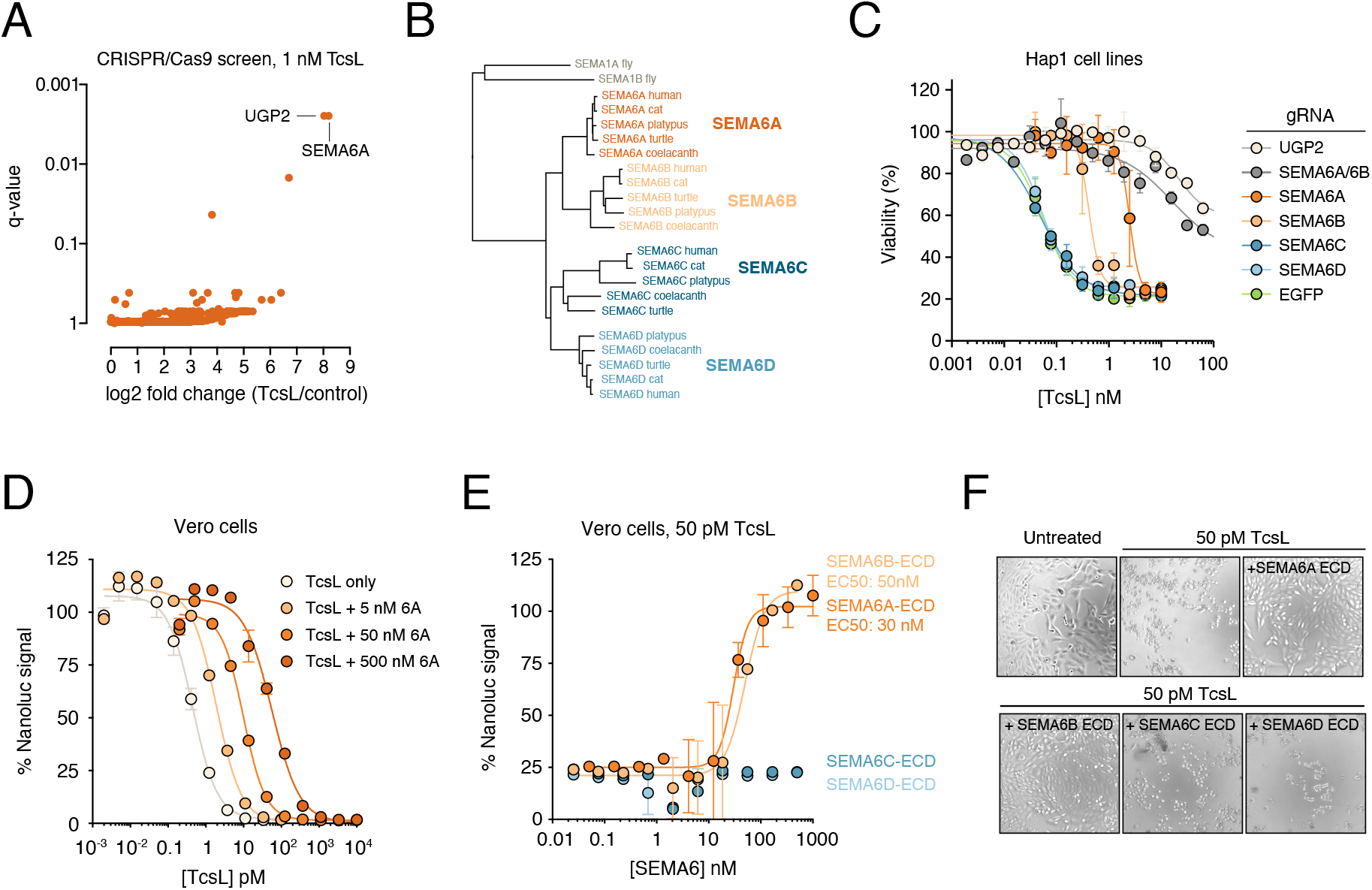
SEMA6A and SEMA6B are host-cell receptors for *C. sordellii* lethal toxin TcsL. (A) Genome-wide CRISPR/Cas9 screen in Hap1 cells identifies factors regulating sensitivity to 1 nM TcsL. (B) Phylogenetic tree of SEMA6 family proteins. (C) Hap1 cells were infected with Cas9 and gRNAs targeting indicated genes and tested for sensitivity to TcsL. (D) Vero cells stably expressing Nanoluciferase viability reporter were treated with increasing amounts of TcsL in the presence of recombinant SEMA6A ectodomain. (E) Vero cells stably expressing Nanoluciferase viability reporter were treated with 50 pM TcsL with increasing amounts of recombinant SEMA6 family ectodomains for 24 hours. (F) Microscopy images of Vero cells treated with TcsL and SEMA6 family ectodomains for 24 hours.

We first validated screen results by knocking out UGP2 and SEMA6A in Hap1 cells. UGP2^KO^ and SEMA6A^KO^ cells were highly resistant to TcsL (fig. S1, C and D). Introduction of wild-type SEMA6A by lentiviral transduction to SEMA6A^KO^ cells rendered the cells more sensitive to the toxin (fig. S1, C and D). SEMA6A is a member of the semaphorin family, which consists of twenty transmembrane and secreted proteins (*14*). The four human SEMA6 class proteins are SEMA6A, SEMA6B, SEMA6C, and SEMA6D (**Fig. 1B**). We generated knockout Hap1 cells with each SEMA6 family member and assayed their sensitivity to TcsL. SEMA6C^KO^ and SEMA6D^KO^ cells were as sensitive to TcsL as control cells (50% growth inhibition GI50: 50 pM; **Fig. 1B**). In contrast, SEMA6B^KO^ cells were approximately 10-fold more resistant (GI50: 500 pM) than control cells, but not as resistant as SEMA6A^KO^ cells (50-fold; GI50: 2.5 nM). Moreover, SEMA6A/SEMA6B double knockout cells were more resistant to TcsL than either knockout alone, suggesting that they act redundantly (**Fig. 1B** and fig. S1D). Consistent with this, HeLa cells that do not express SEMA6A or SEMA6B were highly resistant to TcsL (fig. S1, B and E). Because SEMA6C and SEMA6D are expressed at low levels in Hap1 and HeLa cells (fig. S1B), we further assessed their role in TcsL intoxication by ectopically expressing all SEMA6s in SEMA6A^KO^ cells by lentiviral infection. In contrast to ectopic SEMA6A and SEMA6B expression, SEMA6C and SEMA6D did not make SEMA6A^KO^ cells more sensitive to the toxin (fig. S1F). Thus, SEMA6A and SEMA6B but not SEMA6C or SEMA6D regulate cellular sensitivity to TcsL. This finding reflects the phylogenetic relationship of SEMA6 proteins: SEMA6A and SEMA6B are more closely related to each other than to either SEMA6C or SEMA6D (**Fig. 1B**).

We then expressed and purified the soluble recombinant extracellular domain (rECD) of SEMA6A and tested its effect on TcsL toxicity on Vero cells, a commonly used cell line for studying toxin function. SEMA6A rECD counteracted TcsL toxicity in a dose-dependent manner (**Fig. 1D**). Notably, SEMA6A rECD had a protective effect only when it was added to the cells before or simultaneously with TcsL (fig. S1G). When cells were pretreated for one hour with TcsL, SEMA6A rECD had no effect on toxicity, suggesting that SEMA6A rECD must act before TcsL binds the cell membrane. We also repeated the competition assay with soluble ectodomains of SEMA6B, SEMA6C, and SEMA6D. Consistent with previous experiments, SEMA6B alleviated TcsL toxicity whereas SEMA6C and SEMA6D had no effect (**Fig. 1E** and **Fig. 1F**). Together, these results strongly suggest that TcsL binds SEMA6A and SEMA6B on the cell surface and this interaction is required for TcsL entry into the cell.

One of the primary targets of TcsL during *C. sordellii* infection is the vascular endothelium of the lung (*15*). Therefore, we used immortalized human lung microvascular cells (HULECs) as a physiologically relevant cell line to study the role of SEMA6A and SEMA6B in TcsL intoxication. HULECs express ~4-fold higher levels of SEMA6A and ~11-fold higher levels of SEMA6B than Hap1 cells (**Fig. 2A**). We assayed the sensitivity of HULECs to four related clostridial toxins: TcdA and TcdB from *C. difficile*, TpeL from *C. perfringens*, and TcsL (**Fig. 2B**). TpeL showed low toxicity and TcdA and TcdB were several orders of magnitude less toxic to HULECs than what is reported for other cell types (*16*, *17*). Remarkably, the GI50 of TcsL was ~50 fM, suggesting that only ~200 toxin molecules/cell are lethal to HULECs. Furthermore, recombinant SEMA6A ectodomain but not SEMA6C ectodomain could block TcsL-induced rounding of HULECs in a dose-dependent manner (**Fig. 2C** and **Fig. 2D**). These results strongly suggest that SEMA6A/SEMA6B are the physiologically relevant TcsL receptors in endothelial cells.

**Figure 2.**
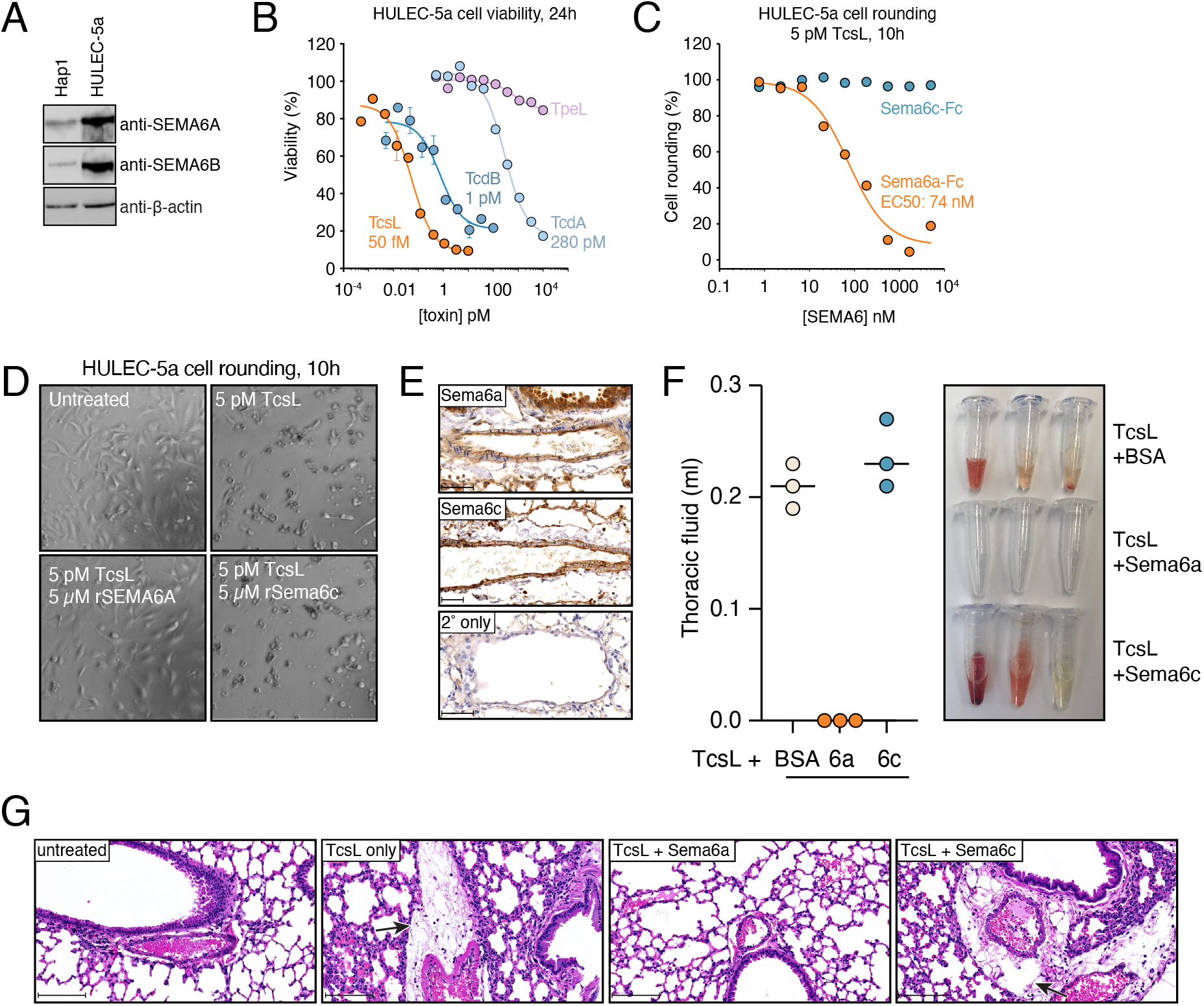
SEMA6A ectodomain protects lung endothelial cells and mouse lungs from TcsL-induced toxicity. A) SEMA6A and SEMA6B protein expression in human lung endothelial cells. (B) HULECs are extremely sensitive to TcsL. HULEC-5a cells were treated with increased amounts of indicated Clostridial toxins. (C) Recombinant mouse Sema6a ectodomain fused to Fc domain protects HULEC-5a cells from TcsL-induced cell rounding. Cells were treated with 5 pM TcsL and increasing amounts of Sema6a and Sema6c ectodomains. (D) Microscopy images of HULEC-5a cells treated with TcsL and recombinant Sema6a and Sema6c ectodomains. (E) Immunohistochemistry images of Sema6a and Sema6b expression in mouse lung tissue sections. (F) Mice were intraperitoneally injected with 15 ng TcsL and 1,000-fold molar excess of Sema6a ectodomain, Sema6c ectodomain or BSA. Thoracic fluid was collected and measured from symptomatic mice four hours after injection. (G) Lung tissue sections of mice treated with indicated conditions. Arrows indicate lung edema induced by TcsL.

We then addressed the role of semaphorins in a mouse model of TcsL intoxication. We first examined the expression of SEMA6A and SEMA6B in mouse lung tissue by immunohistochemistry. Both proteins were highly expressed in lung endothelium and pneumocytes (**Fig. 2E**). We injected mice intraperitoneally with a lethal dose of TcsL together with mouse Sema6a ectodomain-Fc fusion, mouse Sema6c ectodomain-Fc fusion, or BSA (n = 3 for each group). Mice co-injected with BSA or Sema6c-Fc rapidly developed symptoms of TcsL intoxication, including decreased mobility and signs of ataxia and dehydration. Within four hours, all symptomatic mice had a buildup of fluid in the lungs (**Fig. 2F**). Histopathologically, symptomatic mice showed edema surrounding pulmonary vessels (**Fig. 2G**). In contrast, SEMA6A-Fc protected the mice from TcsL-induced symptoms. Mice co-injected with SEMA6A-Fc had no pleural effusion and did not show signs of edema after four hours (**Fig. 2E** and **2F**). Taken together, these data support that SEMA6A, likely acting together with SEMA6B, is the physiologically relevant receptor for TcsL *in vivo*.

We next structurally characterized the interaction between TcsL and SEMA6A by cryo-EM using a shortened TcsL_1285-1804_ construct, which bound recombinant SEMA6A ectodomain with nanomolar apparent affinity (**Fig. 3A** and fig. S2A). Glutaraldehyde cross-linking was used to prevent dissociation of the TcsL_1285-1804_-SEMA6A complex during cryo-EM grid preparation, as described previously (*18–20*). Initial analysis of the dataset revealed that approximately 50% of the SEMA6A dimers were bound to TcsL using this sample preparation approach. Consequently, 2D and 3D classification resulted in two homogenous datasets corresponding to the TcsL-SEMA6A complex (155,353 particle images (fig. S2)) and unliganded SEMA6A (229,275 particle images (fig. S3)), which were used to produce final density maps to overall resolutions of 3.3 and 3.1 Å, respectively (table S2). Local resolution of the TcsL-SEMA6A map ranged from 3.0 Å at the binding interface to >30 Å at the C terminus of TcsL (fig. S2, D and E). Indeed, 3D variability analysis (*21*) revealed continuous motion of the C and N termini of TcsL in contrast to a largely rigid SEMA6A dimer and TcsL-SEMA6A interface (movie S1). A molecular model was built for the well-resolved central domain of TcsL (residues 1401 to 1615), corresponding to approximately 40% of the TcsL construct (**Fig. 3A** and fig. S4).

**Figure 3.**
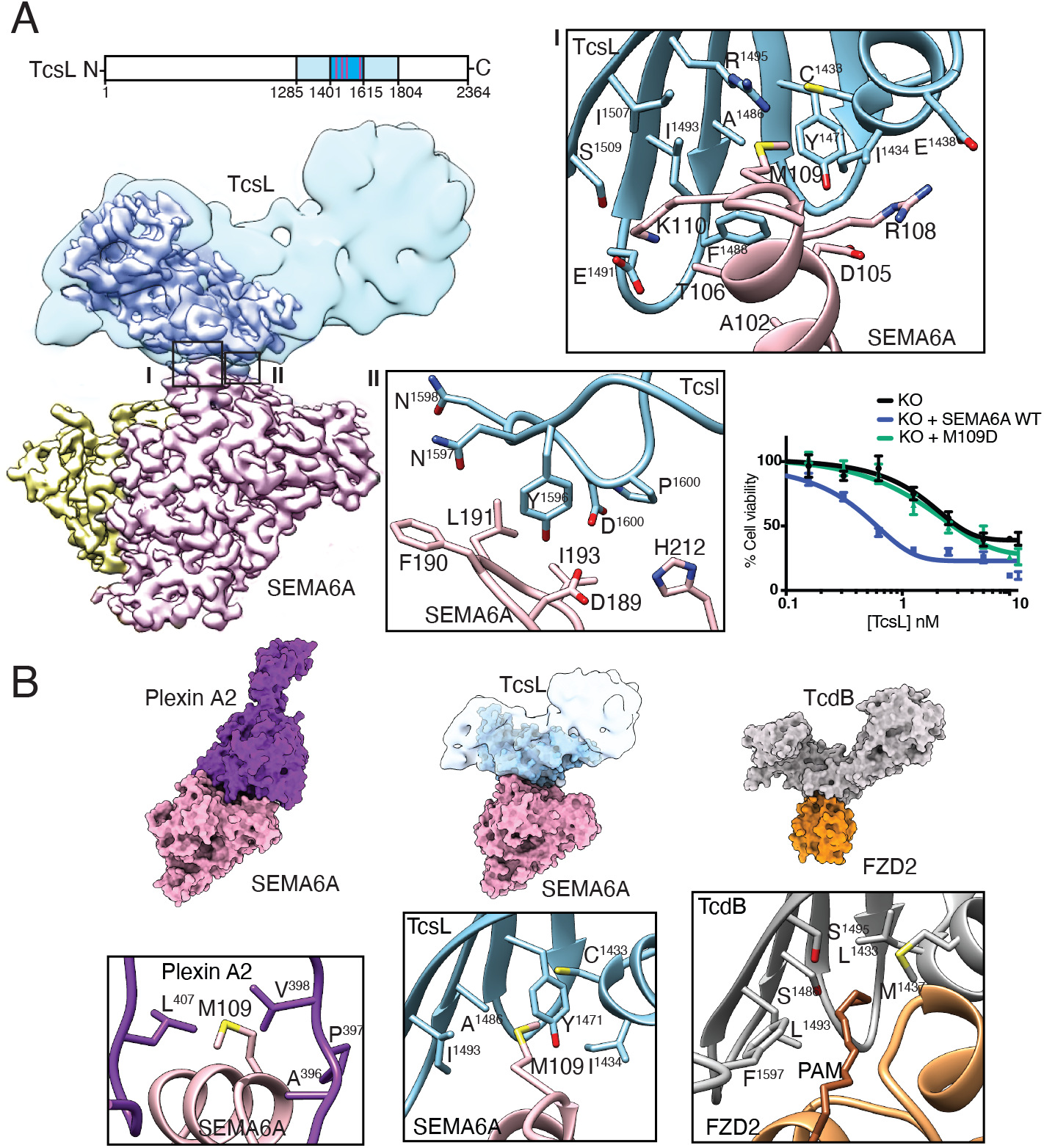
Cryo-EM structure of the TcsL-SEMA6A complex uncovers a receptor-binding region utilized across clostridial toxins. (A) Upper left panel: amino acid sequence of TcsL. White bar – full length TcsL protein, light blue bar – TcsL_1285-1804_ construct used in cryo-EM analysis, dark blue bar – TcsL domain resolved to medium and high resolution (<7 Å) in cryoEM studies. Purple stripes denote residues contacting SEMA6A. Lower left panel: composite cryo-EM map of the Tcsl-SEMA6A complex. SEMA6A monomers are colored pink and yellow, TcsL domain resolved to medium and high resolution (<7 Å) is colored dark blue. Low-pass filtered density (10 Å) of the TcsL protein is shown in light blue. Insets I and II: contact residues between SEMA6A (pink) and TcsL (blue). Lower right panel: validation of SEMA6A M109 as a critical interacting residue with TcsL. Hap1 cells expressing SEMA6A M109D mutant are as resistant to TcsL as SEMA6A KO cells. (B) Comparison of Plexin A2/SEMA6A (PDB: 3OKY) (*25*), TcsL/SEMA6A and TcdB/FZD2 (PDB:6C0B) (*8*) binding interactions. M109 of SEMA6A interacts with hydrophobic residues of Plexin A2, including L407 and V398 (lower left panel). TcsL buries M109 of SEMA6A in a binding pocket containing several hydrophobic residues (lower middle panel). TcdB utilizes a similar binding pocket to interact with the palmitoleic acid moiety of FZD2 (lower right panel).

The architecture of TcsL is similar to previously-determined structures of other clostridial toxins (*8*, *22–24*) (fig. S5). Similarly, unliganded and TcsL-bound SEMA6A structures are in remarkable agreement (fig. S6), indicating that TcsL binding is not accompanied by significant conformational changes in the receptor and that glutaraldehyde treatment had a negligible effect on the SEMA6A structure. The TcsL-SEMA6A interface is formed by discontinuous elements in both proteins that include TcsL residues in an α-helix (1433-38), β-strands and turns (1484-95 and 1505-11) and a flexible loop (1596-1601), and SEMA6A residues in an α-helix (101-108) and in flexible loops (190-193 and 212) (**Fig. 3A** and tables S3 and S4). Several SEMA6A residues (R108, I193, H212) that engage in contacts with TcsL are identical in SEMA6B but not conserved in SEMA6C and D (fig. S7), likely explaining the specificity of TcsL for SEMA6A/B. Critical to the interface is M109 of SEMA6A, which is buried in a TcsL hydrophobic pocket formed by C1433, I1434, Y1471, A1486 and I1493. Indeed, expression of a SEMA6A M109D in SEMA6A knockout cells does not make the cells more sensitive to TcsL, in contrast to wild-type SEMA6A (**Fig. 3A**). TcsL interacts with the SEMA6A receptor at a position partially overlapping the functional site used by the Plexin A2 cognate ligand (**Fig. 3B**)(*25*, *26*). Strikingly, the site on TcsL mediating the SEMA6A interaction is in the same location as the previously reported TcdB-FZD2 binding interface (*8*) (**Fig. 3B** and fig. S8). In fact, the TcsL hydrophobic pocket burying M109 of SEMA6A is analogous to the TcdB hydrophobic pocket burying the FZD2 palmitoleic acid moiety (**Fig. 3B**).

Thus, TcsL and TcdB have evolved to bind different host receptors through the same interaction surface. This surface is highly diverged between the two toxins: only three of the 23 SEMA6A-interacting residues in TcsL are conserved in TcdB, and conversely, four of the 22 residues in TcdB that interact with FZD2 are conserved in TcsL. In stark contrast, 76% of non-interacting surface residues in TcdB are conserved in TcsL (p < 0.0001, Fisher’s exact test), indicating that the divergence is specific to the receptor-binding surface. This surface may represent a more general receptor-binding site in clostridial toxins, as the divergence extends to all members of the family (fig. S9). In particular, the region surrounding the hydrophobic pocket for SEMA6A M109 in TcsL and for the palmitoleic acid moiety in the FZD2/TcdB complex is highly variable between LCTs (**Fig. 4A** and fig. S9).

**Figure 4.**
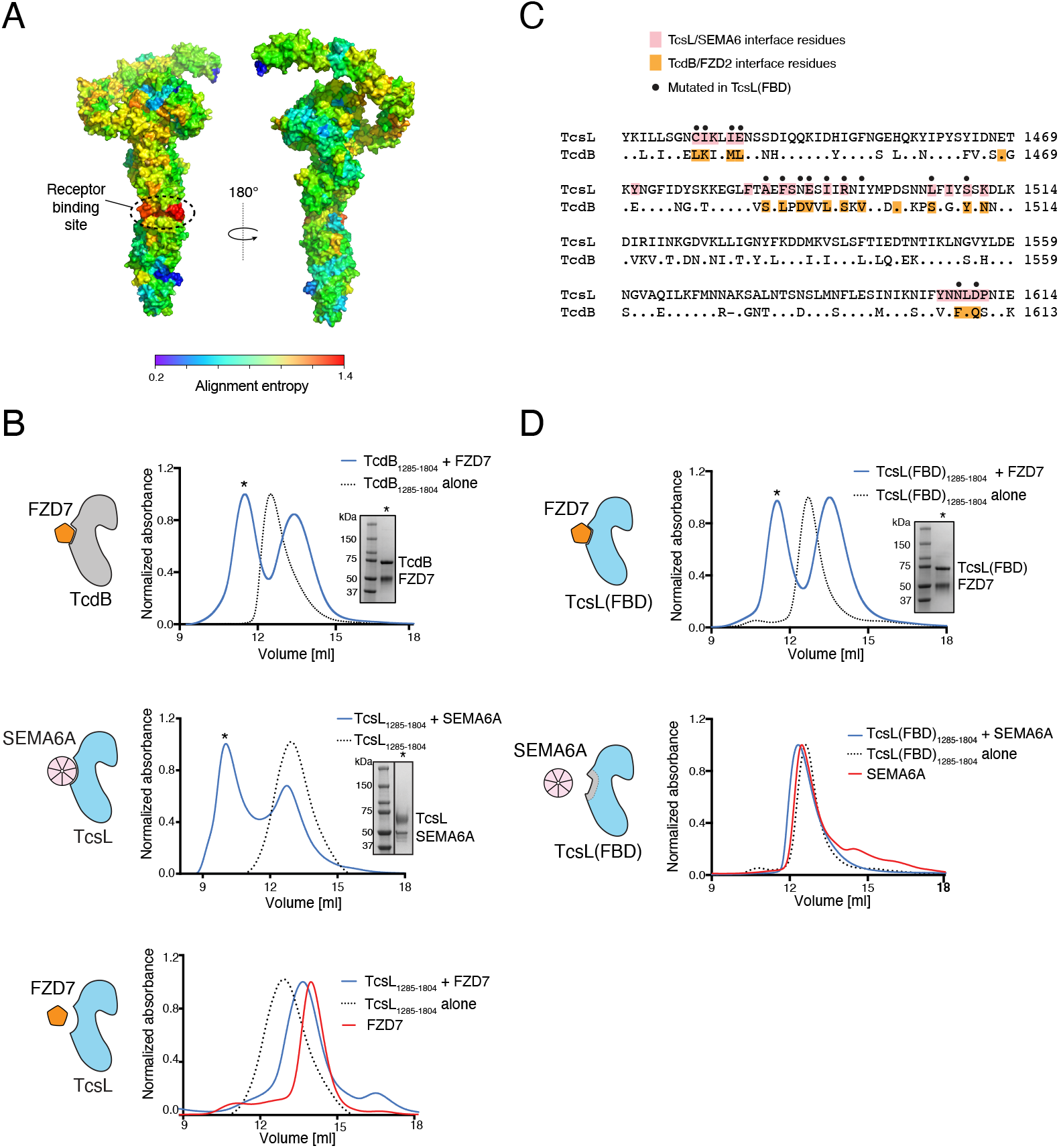
TcsL and TcdB bind different host receptors through the same interface region. (A) Sequence alignment entropy in large clostridial toxin family shown as a rainbow spectrum on the TcdB full-length cryo-EM structure (PDB ID: 6OQ5). Entropy was calculated as a 10-aa moving window. The receptor-binding surface is indicated. (B) SEC profiles and SDS-PAGE analysis of FZD7/TcdB_1285-1804_ (top panel), SEMA6A/TcsL_1285-1804_ (middle panel), and FZD7/TcsL_1285-1804_ (bottom panel). SEC fractions used for SDS-PAGE analysis are highlighted with an asterisk. (C) Sequence alignment between TcsL and TcsB. TcsL residues interacting with SEMA6A are highlighted in pink and TcdB residues contacting FZD2 are highlighted in orange. Black dots denote the 15 mutations introduced in TcsL(FBD)_1285-1804_ variant that resulted in shifting the TcsL binding specificity from SEMA6A to FZD7. (D) SEC profiles and SDS-PAGE analysis of FZD7/TcsL(FBD)_1285-1804_ (TcsL variant with a TcdB-like binding interface; top panel) and SEMA6A/TcsL(FBD)_1285-1804_ (bottom panel). SEC fractions used for SDS-PAGE analysis are highlighted with an asterisk.

We directly tested the role of the evolutionarily divergent surface in receptor specificity with recombinant proteins and size exclusion chromatography. As expected, TcsL_1285-1804_ interacted with SEMA6A, and TcdB_1285-1804_ formed a stable complex with FZD7 (**Fig. 4B**). However, TcsL did not interact with FZD7, consistent with the extensive divergence of the receptor-binding interface (**Fig. 4B**). We then generated a TcsL(FBD)_1285-1804_ variant that had a TcdB-like interface by changing 15 TcsL residues in the receptor-binding interface to those in TcdB (**Fig. 4C** and fig. S8 and S9). Remarkably, these mutations were sufficient to switch TcsL binding specificity: the TcsL_1285-1804_(FBD) hybrid protein robustly interacted with FZD7 but no longer bound SEMA6A (**Fig. 4D**).

These results establish that *C. sordellii* TcsL and *C. difficile* TcdB, two highly similar virulence factors, use the same surface to interact with distinct cognate receptors. The striking evolutionary divergence of this surface among all clostridial toxins suggests that it provides a common, malleable interface that can be modulated to bind structurally unrelated host receptors, allowing the pathogens to adapt to and attack novel host organisms and tissues. Moreover, extending the similarities between toxins, both TcdB and TcsL are likely to interfere with the host receptor function. TcdB, which binds Frizzled, can inhibit Wnt signaling in colonic organoids (*7*). SEMA6A binds TcsL and its endogenous ligand Plexin A2 with overlapping interfaces, suggesting that TcsL could interfere with the semaphorin-plexin signaling axis in vascular endothelium. Further work will address the intriguing possibility that TcsL and other LCTs use their cognate receptor both for gaining entry into the cell and for modulating host cell responses to their advantage.

We cannot exclude the possibility that, like other clostridial toxins (*7*, *10*–*12*), TcsL also binds additional receptors on host cells. However, we did not identify additional factors in CRISPR/Cas9 screens in cells that do not express SEMA6A or SEMA6B. Moreover, if they exist, these receptors must be distinct from those of other LCTs. Our results with physiologically relevant models indicate that SEMA6A and SEMA6B are the major TcsL receptors *in vivo*. Consequently, targeting this interaction with antibodies or small molecules could provide a highly needed therapeutic intervention against rapidly lethal and currently untreatable *C. sordellii* infections.

## Acknowledgements

The authors would like to thank Dr. Amy Tong and Mr. Christopher Mogg for their help with this project.

## Author contributions

Conceptualization, methodology: HL, GB, IK, DS, JPJ, RAM, MT, Investigation: HL, GB, IK, MT, DS, Writing: MT, JPJ, RAM, DS, IK, HL, GB, Resources: SR, HC, JLR, Supervision: JPJ, RAM, MT Funding acquisition: JPJ, RAM, MT,

## Data and material availability

The cryo-EM structure of TcsL/SEMA6A complex will be deposited to PDB and EMDB upon acceptance.

## Supplementary Materials

Materials and Methods

Table S1-S4

Fig S1-S9

Movie S1

## Materials and Methods

### Cell lines

Vero cells were purchased from ATCC (Vero CCL-81) and cultured in complete Dulbecco’s Modified Eagle’s Media (10% FBS, 1 % penicillin/streptomycin), and were transduced by lentivirus expressing nanoluciferase-PEST (Promega). Stable clones were selected by puromycin and limiting dilution. These cells are referred to as Vero-NlucP. Hulec-5a human lung microvascular endothelial cells were purchased from ATCC (CRL-3244) and cultured in Vascular Cell Basal Medium (ATCC PCS-100-030) with the Endothelial Cell Growth Kit-VEGF (ATCC PCS-100-041).

### Genome-wide CRISPR screen

50 million Hap1 cells were infected with TKO V3 library at 200-fold library coverage at multiplicity of infection (MOI) of 0.3. The infected cells were selected with 2 μg/ml puromycin for 3 days (T0) and maintained at a minimum of 100-fold gRNA library coverage at any time point. Cells were passaged until the fourth day (T4) to allow sufficient time for protein turnover. 7 million cells were seeded into two 10cm plates per condition and allowed to recover overnight after trypsinization. At T5, TcsL was added at a final concentration of 0.1nM or 1nM, and the cells were further incubated for 48 hours. At T7, cells were washed with 1xPBS and allowed to repopulate in normal growth media (IMDM/10% FBS). The untreated population of cells (negative controls) were passaged every three days in parallel. The medium of toxin treated cells were replenished every 3 days. Once surviving colonies were visible, cells were trypsinized and re-seeded to facilitate repopulation. Once the toxin treated cells reached 100% confluency, the cells were collected with the corresponding untreated population. Genomic DNA was extracted from the frozen pellets using QIAamp DNA Blood Maxi Kit (Qiagen) following the manufacturer’s protocol.

### Next-generation sequencing library preparation

The gRNA sequences were PCR amplified from the extracted genomic DNA. Each amplified sample was then barcoded and processed on Illumina Next-seq high-output mode at a read depth of at least 5 million reads per sample. MAGeCK software (*27*) was used to generate rankings for positively enriched genes.

### CRISPR screen validation

The gRNAs targeting SEMA6A, SEMA6B, SEMA6C, SEMA6D and UGP2 were chosen from TKOv3 library and were cloned into pX459 (Addgene plasmid #62988; gift from Feng Zhang) and lentiCRISPRv2 (Addgene plasmid #52961; gift from Feng Zhang). Plasmids were transfected into HAP1 cells seeded on a 6-well plate with Turbofectin (OriGene) following the manufacturer’s protocol. One day post-transfection, the medium was changed to medium containing 2 μg/ml puromycin and was further selected for 3 days. Cells were washed and moved to a 10 cm plate with fresh growth medium with no antibiotics.

Wild-type and knock-out cells were seeded on a 96-well plate a day before toxin application at <40% confluency. Toxins were serially diluted in 1xPBS with 10% glycerol before applying to cells. Cells were incubated with toxins for 24 to 48 hours. Cell viability was measured either using AlamarBlue dye (Invitrogen) or CellTiter-Glo reagent (Promega) following the manufacturer’s protocol.

For re-expression of SEMA6A, SEMA6B, SEMA6C, and SEMA6D, full-length genes were cloned into a pLenti6.2 plasmid with a C-terminal FLAG and V5 tags. Lentiviruses were packaged by transfecting the lentiviral plasmid with VSV-G and psPAX packaging plasmids into HEK293T cells. Virus-containing media was collected three days post-transfection. The media containing packaged virus was added to HAP1-SEMA6A knock-out cells in a 1:10 ratio and selected on 10 μg/ml blasticidin for 7 to 10 days. The expression of the target construct was confirmed by SDS-PAGE and Western blotting with anti-FLAG-HRP antibody (Sigma).

### Site-directed mutagenesis

Mutations were generated with Quikchange site-directed mutagenesis (Agilent) following manufacturer’s protocol.

### Vero cell experiments

Vero-NlucP cells were plated at 4000 cells/well in a white-walled clear bottom 96-well plate (Corning) and incubated overnight to attach. For toxicity experiments, TcsL was added at the indicated concentrations and incubated for 24 or 48 hours (indicated in the figures). Nanoluciferase levels were measured using the Nano-Glo Luciferase Assay System (Promega) as per the manufacturer’s instructions. For SEMA6 competition experiments, SEMA6 proteins were added immediately before TcsL at the indicated concentrations and the assay proceeded as above.

### Hulec-5a cells

Hulec-5a cells were plated in white-walled clear-bottom 96-well plates at 4000 cells/well and left to attach overnight. TcsL and/or SEMA6 proteins were added as indicated. For cell viability experiments, cells were incubated for 48 hrs at 37°C, 5% CO2 and cell viability was measured using CellTiter-Glo (Promega) as per the manufacturer’s instructions. For cell rounding experiments, media was replaced by complete media containing 1μM CellTracker Orange CMRA (Molecular Probes). After 60 mins, excess dye was removed by media exchange with complete media. Cells were then incubated with 5 pM TcsL and various concentrations of SEMA6 proteins as indicated. Cells were incubated for 10 hrs before imaging. CellTracker-labeled cells were evaluated on a Cellomics ArrayScan VTI HCS reader (Thermo Scientific) using the target acquisition mode, a 10x objective, and a sample rate of 150 objects per well. After recording all image data, the cell rounding and shrinking effects of TcsL intoxication were calculated using the cell rounding index (CRI), a combined measure of the length-to-width ratio (LWR) and area parameters. Dose response curves were plotted and fit to a sigmoidal function (variable slope) to determine EC_50_ using Prism software (GraphPad Software).

### Recombinant proteins

Recombinant human SEMA6B, SEMA6C, and SEMA6D ectodomains fused to Fc domain were purchased from R&D Systems. Mouse Sema6a-Fc and Sema6c-Fc expression constructs were a gift from Woj Wojtowicz (Addgene plasmids 72163 and 72167, respectively). Plasmid pHis1522 encoding his-tagged TcsL was synthesized and codon optimized for *Bacillus megaterium* (Genscript). To express and isolate recombinant TcsL, transformed *B. megaterium* was inoculated into LB containing tetracycline and grown to an A600 of 0.7, followed by overnight xylose induction at 37 °C. Bacterial pellets were collected, resuspended with 20 mM Tris (pH 8)/0.5 M NaCl, and passed twice through an EmulsiFlex C3 microfluidizer (Avestin) at 15,000 psi, then clarified by centrifuging for 18,000 x g for 20 min. TcsL was purified by nickel affinity chromatography followed by anion exchange chromatography using HisTrap FF and HiTrap Q columns (GE Healthcare), respectively. Fractions containing TcsL were verified by SDS-PAGE, then pooled and diafiltered with a 100,000 MWCO ultrafiltration device (Corning) into 20 mM Tris (pH 7.5)/150 mM NaCl. Finally, glycerol was added to 15% v/v, the protein concentration was estimated by A280 (using ext. coefficient 300205), and stored at −80°C.

TcsL fragments (1285-1804) and all mutants thereof were synthesized (IDT) and cloned into pET Champion vector with an N-terminal 6xHIS-SUMO-Strep-TEV sequence and a C-terminal TEV-6xHIS. Positive clones were verified by sequencing. NiCo21(DE3) competent E. coli (NEB) were transformed and inoculated in LB media with kanamycin and grown to an A600 of 0.6. Protein expression was induced by the addition of 1 mM IPTG for 4 hours at 23 °C. Cells were pelleted and protein was purified similarly to TcsL, except after nickel affinity chromatography, they were passed through a size exclusion column and eluted into 20mM Tris pH 8, 150 mM NaCl.

Mouse SEMA6A-Fc and SEMA6c-Fc expression constructs were a gift from Woj Wojtowicz (Addgene plasmids 72163 and 72167, respectively) and human SEMA6A was gene-synthesized (GeneArt). All SEMA6 constructs were transiently expressed in suspension HEK 293F cells. Proteins were purified by either Protein A affinity or by using ion exchange chromatography (MonoQ 10/100 GL column, GE Healthcare) with a 0 mM–1 M NaCl gradient. These steps were followed by gel filtration chromatography (Superdex 200 Increase, GE Healthcare) in 20 mM Tris pH 9.0 and 150 mM NaCl buffer.

Recombinant FZD7 was expressed and purified as previously described (*28*). Briefly, a human FZD7 (residues 42-179)-mVenus construct was transiently expressed in suspension HEK 293F cells and purified using Ni-NTA affinity chromatography. The protein was eluted with an increasing gradient of imidazole with a maximum concentration of 500 mM, in a buffer containing 20 mM Tris, pH 8.0, 500 mM NaCl and 5% (v/v) glycerol. This step was followed by gel filtration chromatography (Superdex 200 Increase, GE Healthcare) in 20 mM Tris pH 8.0 and 150 mM NaCl buffer.

To test the formation of toxin-receptor complexes, TcsL_1285-1804_ and TcsL(FBD)_1285-1804_ and TcdB_1285-1804_ (positive control) were mixed with 5-fold molar excess of SEMA6A or FZD7 and incubated at room temperature for 30 min. This was followed by gel filtration chromatography in 20 mM Tris pH 9.0, 150 mM NaCl buffer.

### Biolayer interferometry (BLI)

To determine the binding kinetics for recombinant His-tagged TcsL_1285-1804_ and human SEMA6A, BLI experiments were performed on an Octet Red96 instrument (FortéBio) at 25 °C. All proteins were diluted in kinetics buffer (PBS, pH 7.4, 0.01% (w/v) BSA, 0.002% (v/v) Tween-20). After 10-30 μg/mL of His-tagged TcsL_1285-1804_ was immobilized on Ni-NTA sensors, the baseline was established for 60 s. Subsequently, the loaded biosensors were dipped into wells containing serial dilutions of human SEMA6A, to determine the rate of association. Sensors were then dipped back into kinetics buffer to establish the dissociation rate. The curves were fitted to a 1:1 binding model and the apparent dissociation constant (KD) was evaluated using FortéBio’s Data Analysis software 9.0. Reported values represent the average of four independent experiments with standard error of the mean (SEM).

### TcsL-SEMA6A complex formation and glutaraldehyde crosslinking

TcsL_1285-1804_ was combined with excess of SEMA6A and purified by gel filtration chromatography (Superdex 200 Increase, GE Healthcare) in 20 mM Hepes pH 7.0 and 50 mM NaCl. Fractions containing the complex were concentrated to 0.12 mg/ml and proteins were crosslinked by addition of 0.05% (v/v) glutaraldehyde (Sigma Aldrich) and incubated at room temperature for 60 min. The reaction was stopped by addition of Tris-HCL pH 7.0 to a final concentration of 50 mM. Subsequently, the complex was concentrated to 0.9 mg/ml, spun down for 30 min at 14,500 x g and directly used for cryo-EM grid preparation.

### Cryo-EM data collection and image processing

Homemade holey gold grids (*29*) were glow discharged in air for 15 s before use. TcsL-SEMA6A (3 μl, 0.9 mg/mL) was applied to grids, blotted for 12 s, and frozen in a mixture of liquid ethane and propane (*30*) using a modified FEI Vitrobot (maintained at 4 °C and 100% humidity). Data collection was performed on a Thermo Fisher Scientific Titan Krios G3 operated at 300 kV with a Falcon 4 camera automated with the EPU software. A nominal magnification of 75,000 (calibrated pixel size of 1.03 Å) and defocus range between 1.6 and 2.2 μm were used for data collection. Exposures were fractionated as movies of 30 frames with an exposure rate of 5 electrons/pixel/second and total exposure of 45.2 electrons/Å^2^.

A total of 4581 raw movies were obtained for the TcsL-SEMA6A sample. Image processing was carried out in cryoSPARC v2 (*18*). Motion correction was performed with Patch Motion algorithm and CTF parameters were estimated from the average of aligned movie frames with Patch CTF. 4,659,284 particle images were selected by template matching and individual particle images were corrected for beam-induced motion using local motion algorithm (*31*) within cryoSPARC v2. Ab initio structure determination and classification revealed that ~50% particle images corresponded to a SEMA6A dimer and the remaining particles to a SEMA6 dimer-TcsL complex. The overwhelming majority of particles for the SEMA6A-TcsL complex had one TcsL molecule bound to the SEMA6A dimer, with no evidence of two TcsL molecules bound to the dimer that could be clearly identified for this cross-linked sample. Multiple rounds of heterogeneous refinement followed by non-uniform refinement resulted in a 3.3 Å resolution map of the SEMA6A-TcsL complex (155,353 particle images) and 3.1 Å resolution map of unliganded SEMA6A (229,275 particle images)

### Model building

An initial model for TcsL was created using the Phyre2 server (*32*) with the TcdB crystal structure (PDB: 6C0B) as a reference. The atomic coordinates of SEMA6A dimer (PDB: 3OKW) were manually fitted into the density map using UCSF Chimera (*33*) to generate a starting model, followed by manual rebuilding using Coot (*34*). All models were refined using the phenix.real_space_refine (*35*) with secondary structure and geometry restraints. The final models were evaluated by MolProbity (*36*). Statistics of the map reconstruction and model refinement are presented in table S2.

### Mouse studies

Female C67/Bl6J mice, 8 weeks of age, were intraperitoneally injected with TcsL together with mouse Sema6a-Fc, mouse Sema6c-Fc, or BSA. In brief, 15 ng of TcsL was mixed with a 1000-fold molar excess of each protein at 4 °C on a rotator for one hour prior to injection. Four hours after intoxication, animals were euthanized and fluid present in the thoracic cavity was collected for analysis. For immunohistochemistry, lung tissue was fixed 4% paraformaldehyde for 48 h, processed through an ethanol series to xylene and then paraffin using a Leica ASP300 automatic tissue processor and embedded in paraffin wax using Leica Histocore Arcadia H. Samples were sectioned at 4.5 μm with a Leica RM2255 semi-automatic microtome. Slides for IHC were dewaxed, rehydrated and sections were treated with 3% H2O2 in PBS to kill endogenous peroxidase activity. Antigen retrieval was performed using a 10 mM Tris/1mM EDTA/0.05% Tween 20 (pH 9) buffer solution in a microwave for 15 minutes. Primary antibody was applied at 4 °C overnight using SEMA6A (R&D, AF1615, goat, 1:100 dilution) and SEMA6B (R&D, AF2094, goat, 1:100 dilution). Rabbit anti-goat secondary (Vector Labs BA-1000) was applied for 30 minutes followed by ABC regent (Vector Labs PK-6100) for 25 minutes and developed with DAB (Vector Labs SK-4100) for 4 minutes or less. Tissue was counterstained with Harris Hematoxylin (HHHS-128, Sigma) for 8 minutes and mounted with Shur Mount (Electron Microscopy Sciences 17991-01). Stained slides were digitized at 40x using a Nanozoomer 2.0 HT (Hamamatsu Photonics) and images of stained samples were analyzed using NPD.view2 software (Hamamatsu U12388-01).

---

**Figure S1.**
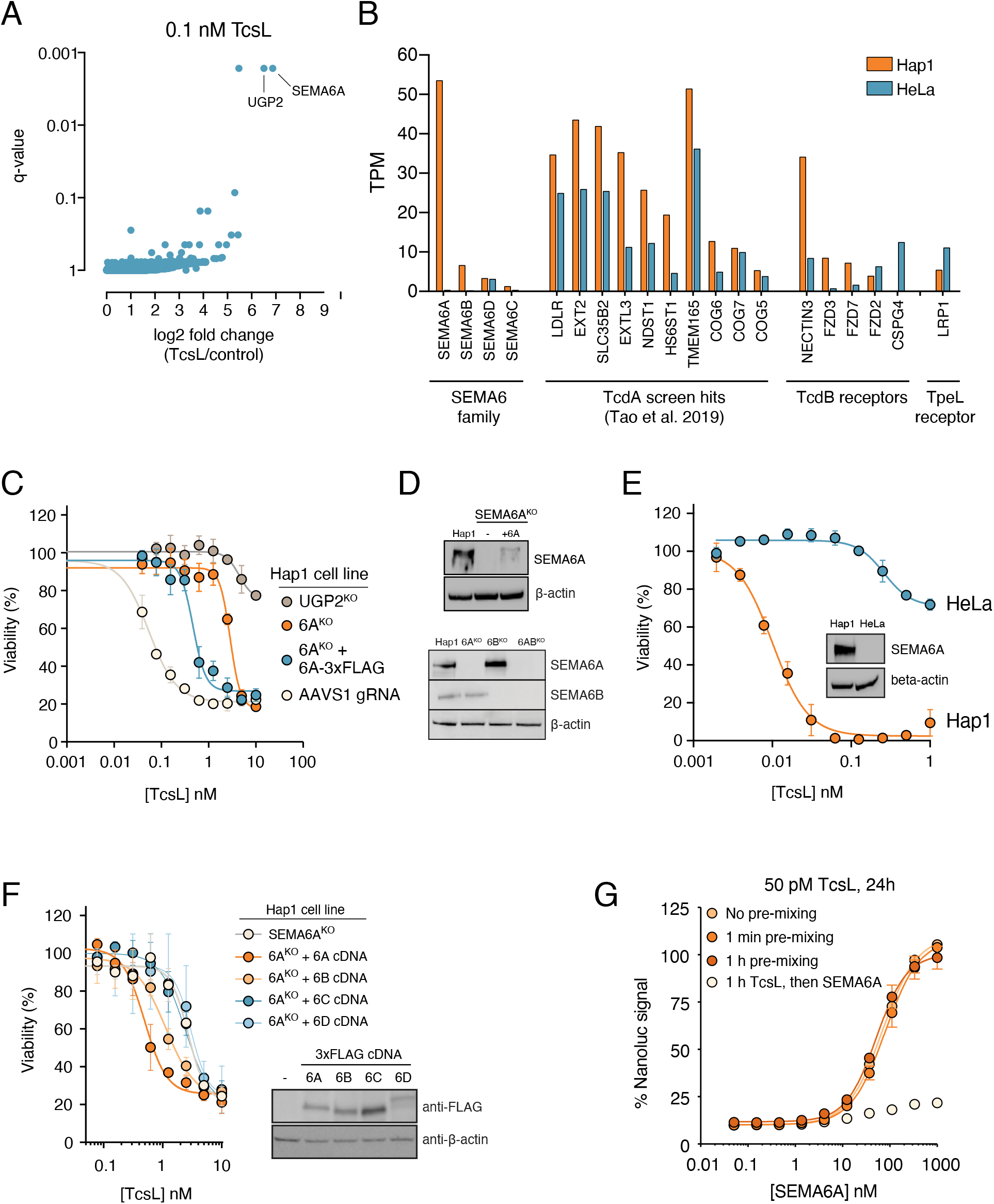
Validation of SEMA6A and SEMA6B as host factors required for TcsL intoxication. (A) Genome-wide CRISPR/Cas9 screen in Hap1 cells identifies factors regulating sensitivity to 0.1 nM TcsL. (B) Expression of SEMA6 family genes and known clostridial toxin receptors and host cell factors in Hap1 and HeLa cells based on Human Protein Atlas (proteinatlas.org). (C) Left, SEMA6A and UGP2 knockout cells were generated with CRISPR/Cas9. SEMA6A-3xFLAG was ectopically expressed in SEMA6A knockout cells by lentiviral infection. (D) Top, SEMA6A expression in wild-type Hap1 cells, SEMA6A^KO^ cells, and in SEMA6A^KO^ cells ectopically expressing SEMA6A-3xFLAG. Bottom, SEMA6A and SEMA6B expression in wild-type Hap1 cells, SEMA6A^KO^ cells, SEMA6B^KO^ cells, and SEMA6A^KO^/SEMA6B^KO^ cells (E) Hap1 and HeLa cells were treated with increasing concentrations of TcsL. Inset, expression of SEMA6A in Hap1 and HeLa cells was assessed by Western blot. (F) Hap1 SEMA6A^KO^ cells were infected with lentiviruses expressing 3xFLAG tagged SEMA6 family proteins, and tested for TcsL sensitivity. Expression of SEMA6 proteins was validated with western blotting (right panel) (G) SEMA6A ectodomain protects Vero cells from TcsL toxicity only when added simultaneously with the toxin. SEMA6A and TcsL were added to Vero cells simultaneously or after 1-minute or 1-hour pre-incubation. Alternatively, TcsL was added for 1 hour prior to treatment with SEMA6A.

**Figure S2.**
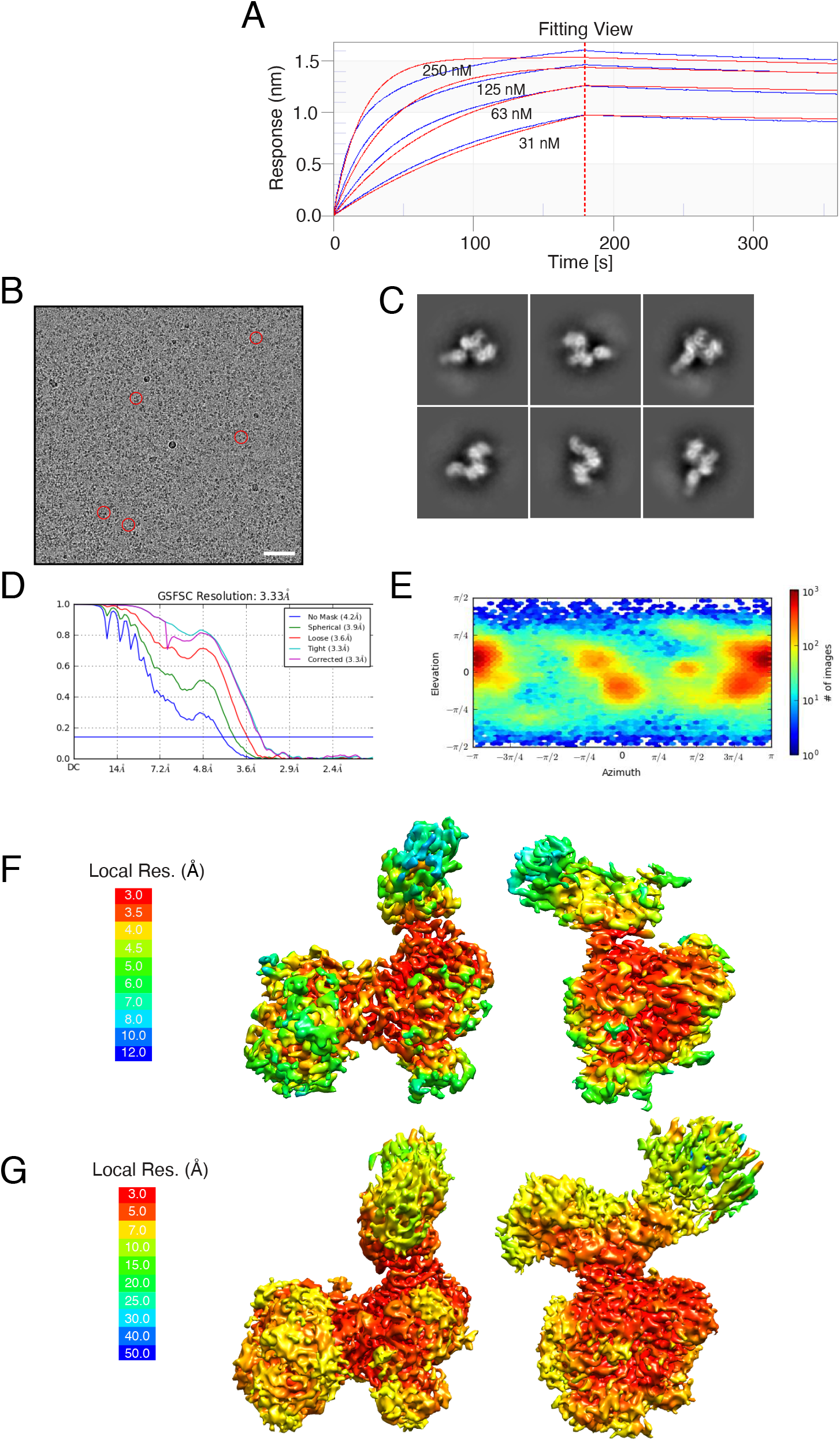
Biochemical and cryo-EM analysis of the SEMA6A/TcsL complex. (**A**) Representative binding curves for the TcsL_1285-1804_/SEMA6A interaction. In this analysis, His-tagged TcsL_1285-1804_ was immobilized on the Ni-NTA biosensor. The average apparent binding affinity from four independent experiments is 2.4 ± 0.8 nM. The data (blue) were fitted using a 1:1 binding model (red). (**B**) An example of cryo-EM micrograph. Scale bar is 50 nm. (**C**) Selected 2D class averages of the TcsL/SEMA6A complex. (**D**) GFFSC curve of the final 3D non-uniform refinement of the TcsL/SEMA6A complex in Cryosparc v2. (**E**) Viewing direction distribution of the TcsL/SEMA6A data. **F** and **G** – local resolution (Å) plotted on the surface of cryo-EM maps countered at different levels to show weak density of the TcsL C terminus.

**Figure S3.**
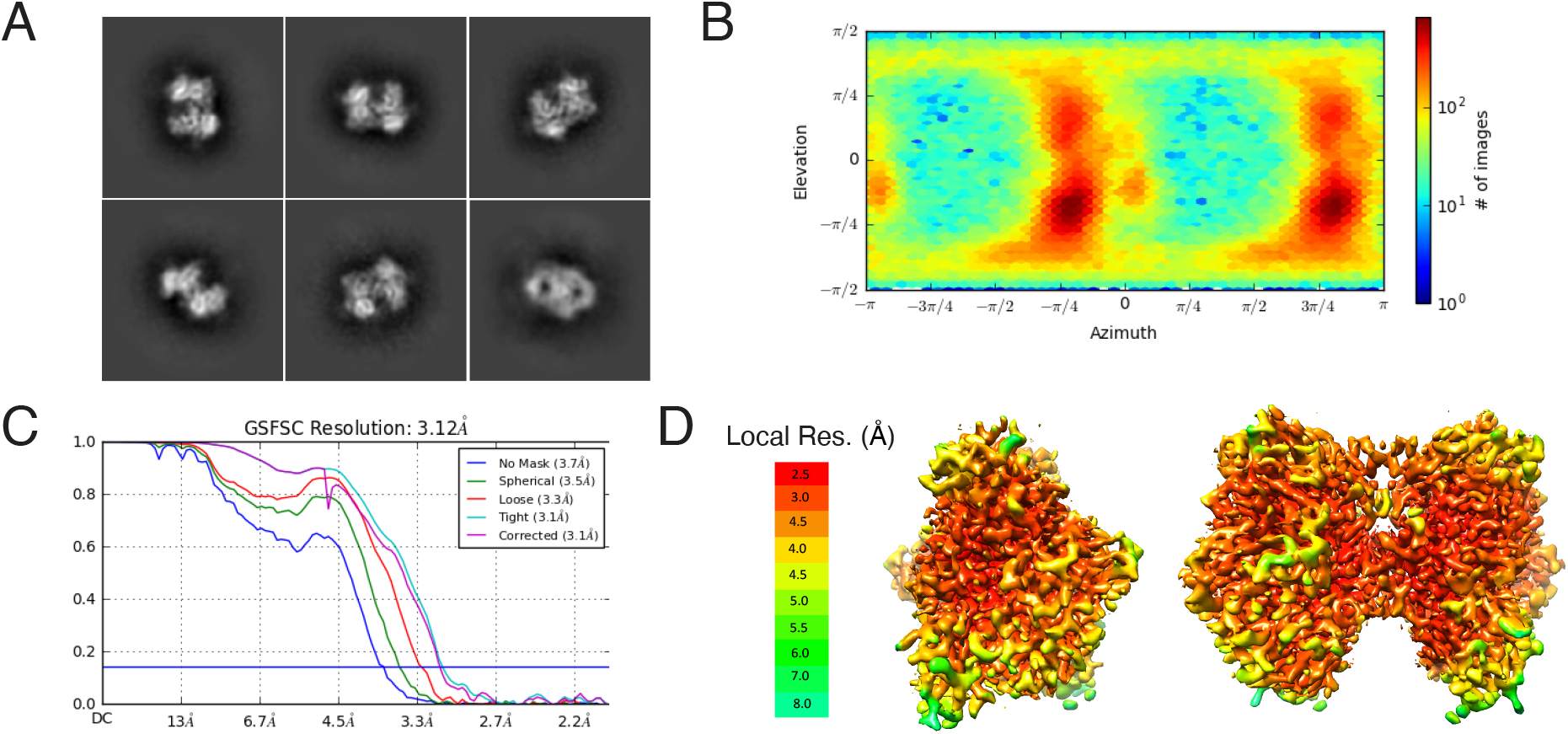
Cryo-EM analysis of SEMA6A. (**A**) Selected 2D class averages of SEMA6A dimer. (**B**) GFFSC curve of the final 3D non-uniform refinement of the SEMA6A dimer in Cryosparc v2. (**C**) Viewing direction distribution of the SEMA6A dimer data. (**D**) Local resolution (Å) plotted on the surface of the SEMA6A cryo-EM map.

**Figure S4.**
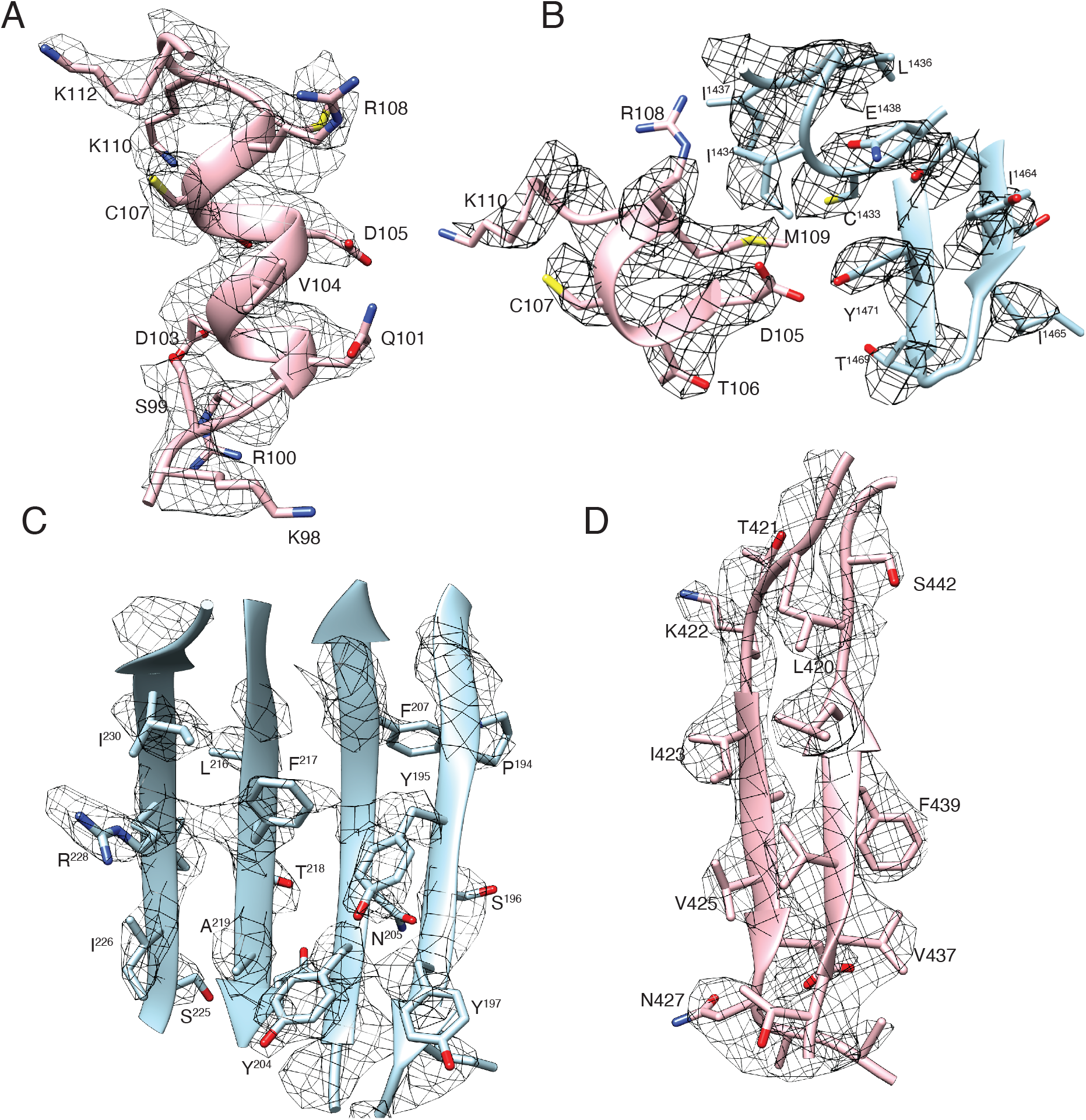
Atomic model built into the SEMA6A/TcsL map. (**A**) α-helix of SEMA6A (residues 102-110); a main interaction site with TcsL. (**B**) Part of the TcsL hydrophobic pocket (blue) accommodating M109 of SEMA6A (pink). (**C**) and (**D**) Examples of β-sheets densities of TcsL (**C**) and SEMA6A (**D**).

**Figure S5.**
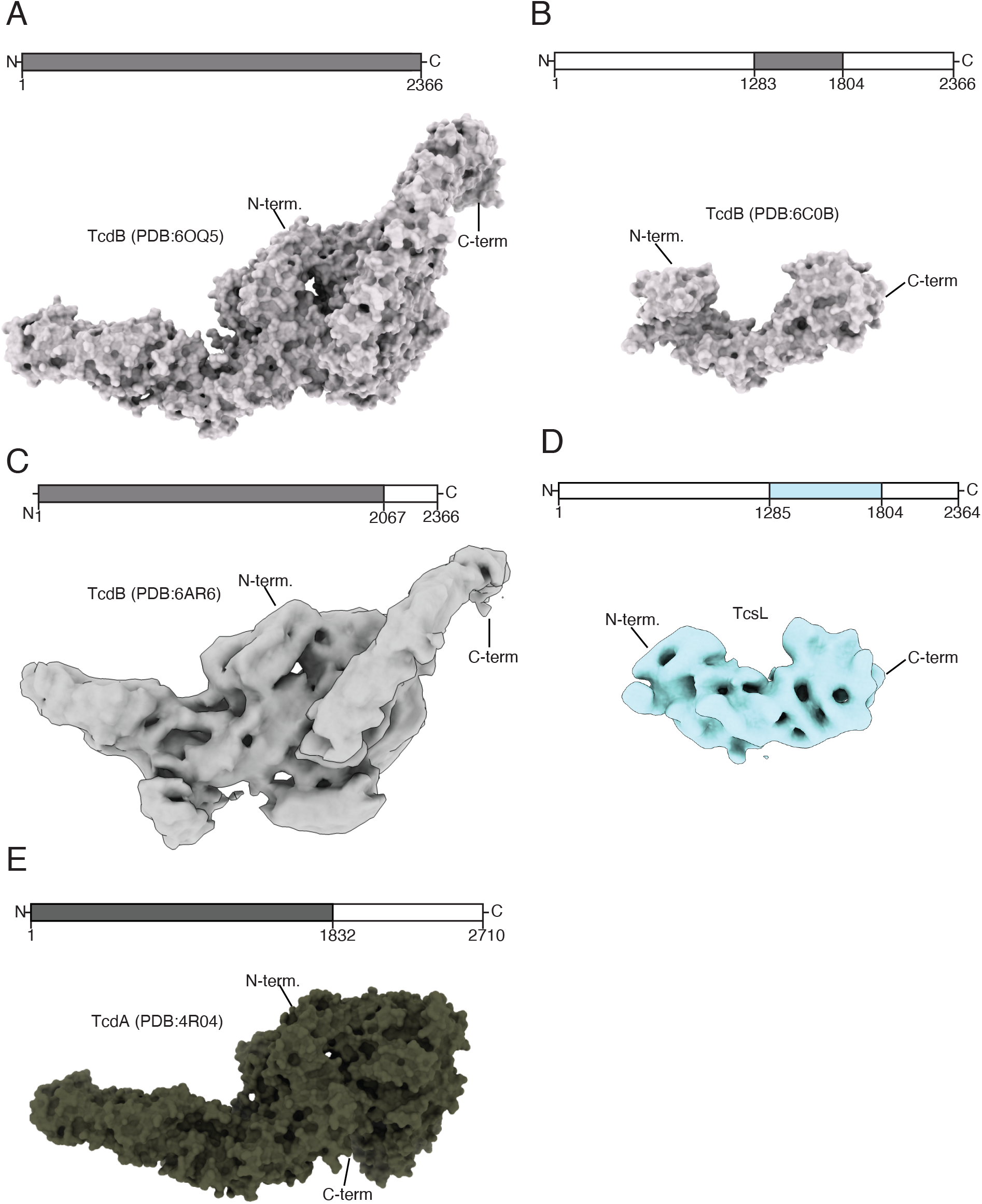
Structural comparison of clostridial toxins. (**A**) X-ray structure of the full-length TcdB toxin (PDB:6OQ5) (*23*). (**B**) X-ray structure of TcdB toxin fragment (residues 1283-1804) (PDB:6C0B) (*8*). (**C**) Cryo-EM structure of TcdB toxin fragment (residues 1-2067) (PDB:6AR6) (*24*). (**D**) Low-pass filtered (10 Å) cryo-EM structure of TcsL toxin fragment (residues 1285-1804). (**E**) X-ray structure of TcdA toxin fragment (residues 1-1832) (PDB:4R04)(*22*).

**Figure S6.**
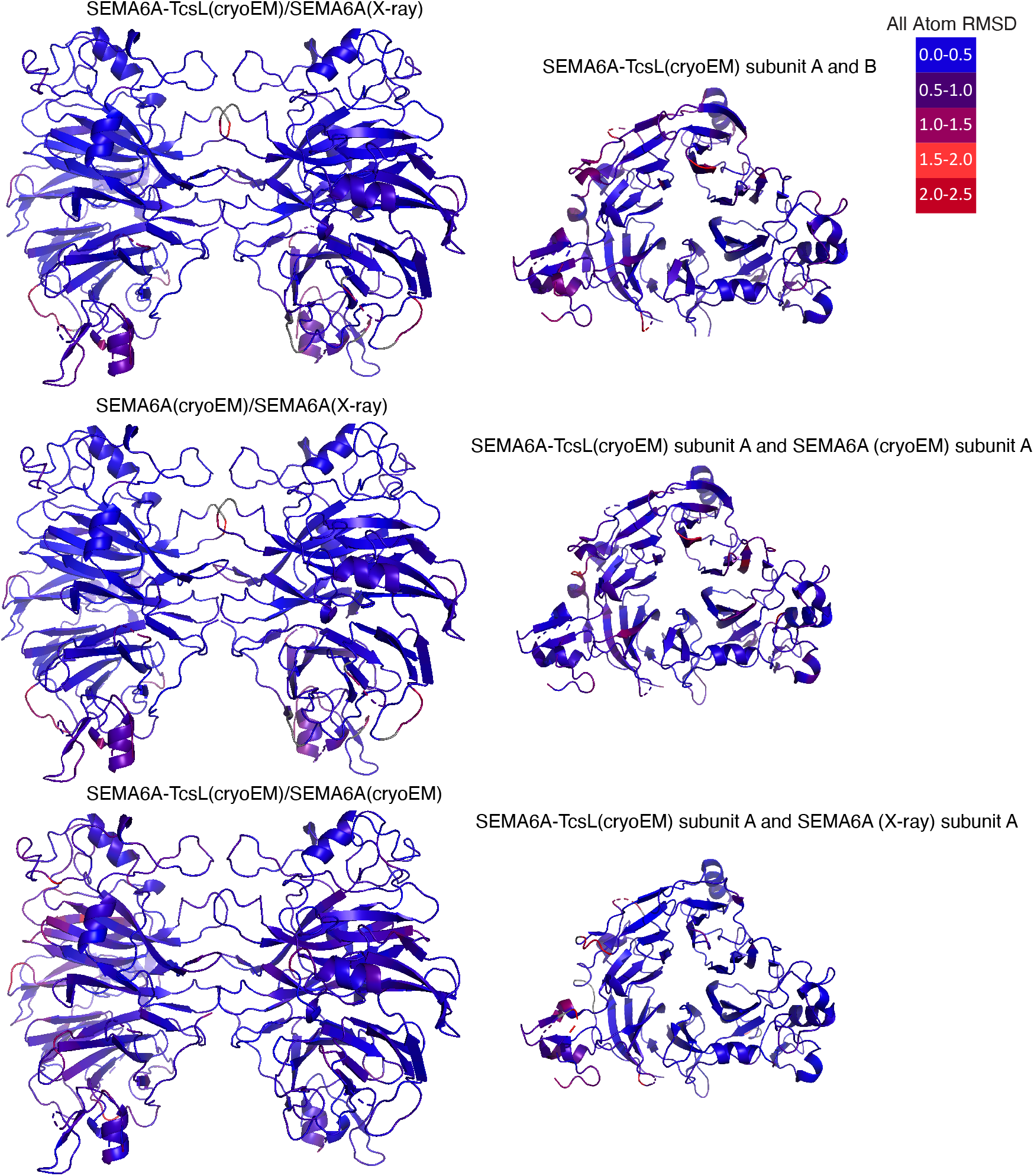
Comparison of cryo-EM SEMA6A structures with SEMA6A X-ray structure (PDB: 3OKW) (*25*). All-atom RMSD values between SEMA6A/TcsL cryo-EM and SEMA6A dimer X-ray structures (upper left panel), SEMA6A cryo-EM and SEMA6A dimer X-ray structures (middle left panel), SEMA6A cryo-EM and SEMA6A/TcsL cryo-EM structures (lower left panel), individual SEMA6A subunits in the SEMA6A/TcsL cryo-EM structure (upper right panel), TcsL-bound SEMA6A subunit in the SEMA6A/TcsL cryo-EM structure and SEMA6A subunit in SEMA6A cryo-EM structure (middle right panel), and TcsL-bound SEMA6A subunit in SEMA6A/TcsL cryo-EM structure and SEMA6A subunit in SEMA6A X-ray structure (lower right panel). All atom RMSD values were calculated using Pymol (*16*) and plotted on the surface of SEMA6A dimers (left) and monomers (right).

**Figure S7.**
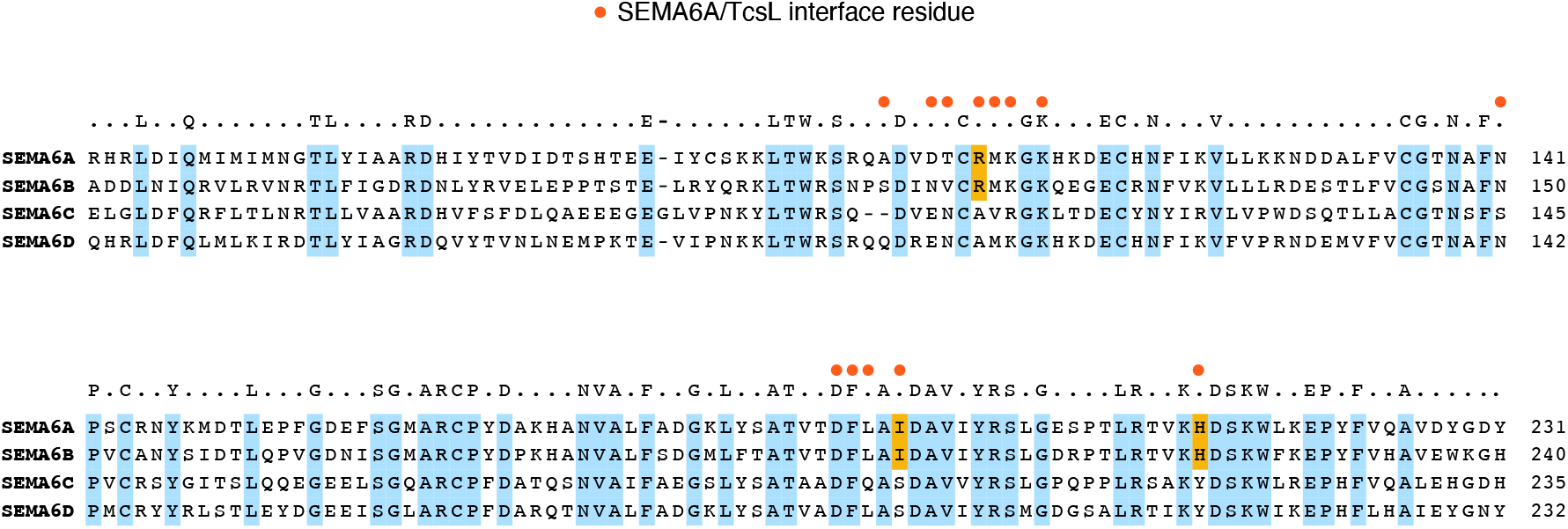
Sequence alignment of SEMA6 family proteins. Residues conserved in all four SEMA6 family proteins are denoted in light blue. SEMA6A residues forming contacts with TcsL are indicated as red circles. Contact residues that differ between SEMA6A/SEMA6B and SEMA6C/SEMA6D are colored orange.

**Figure S8.**
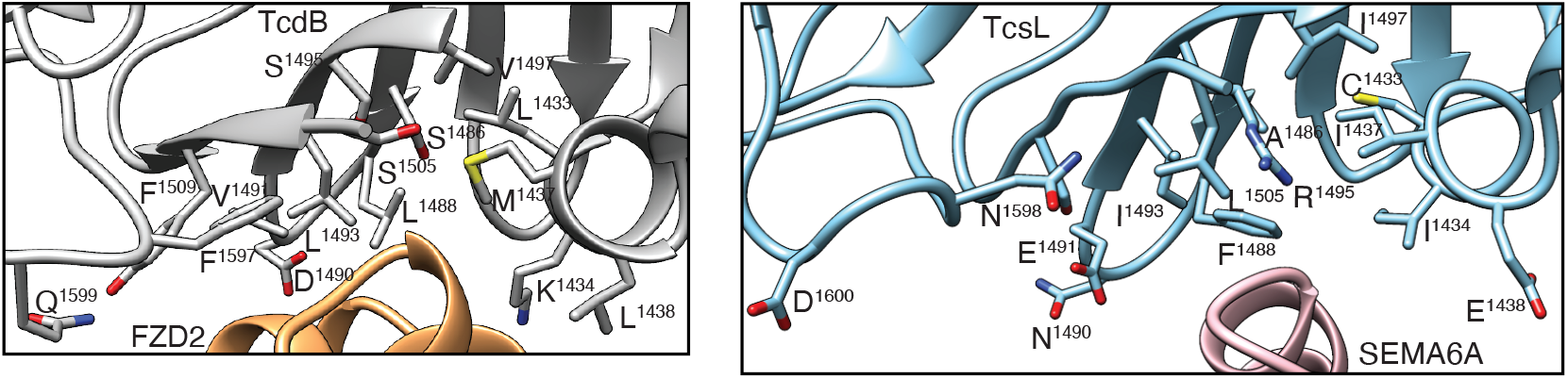
Structural details for receptor-binding region utilized by TcsL and TcdB clostridial toxins. Positions of introduced point mutations in TcsL(FBD)_1285-1804_ are highlighted in the structures of TcdB/FZD2 (PDB:6C0B) (left panel) and TcsL/SEMA6A (right panel).

**Figure S9.**
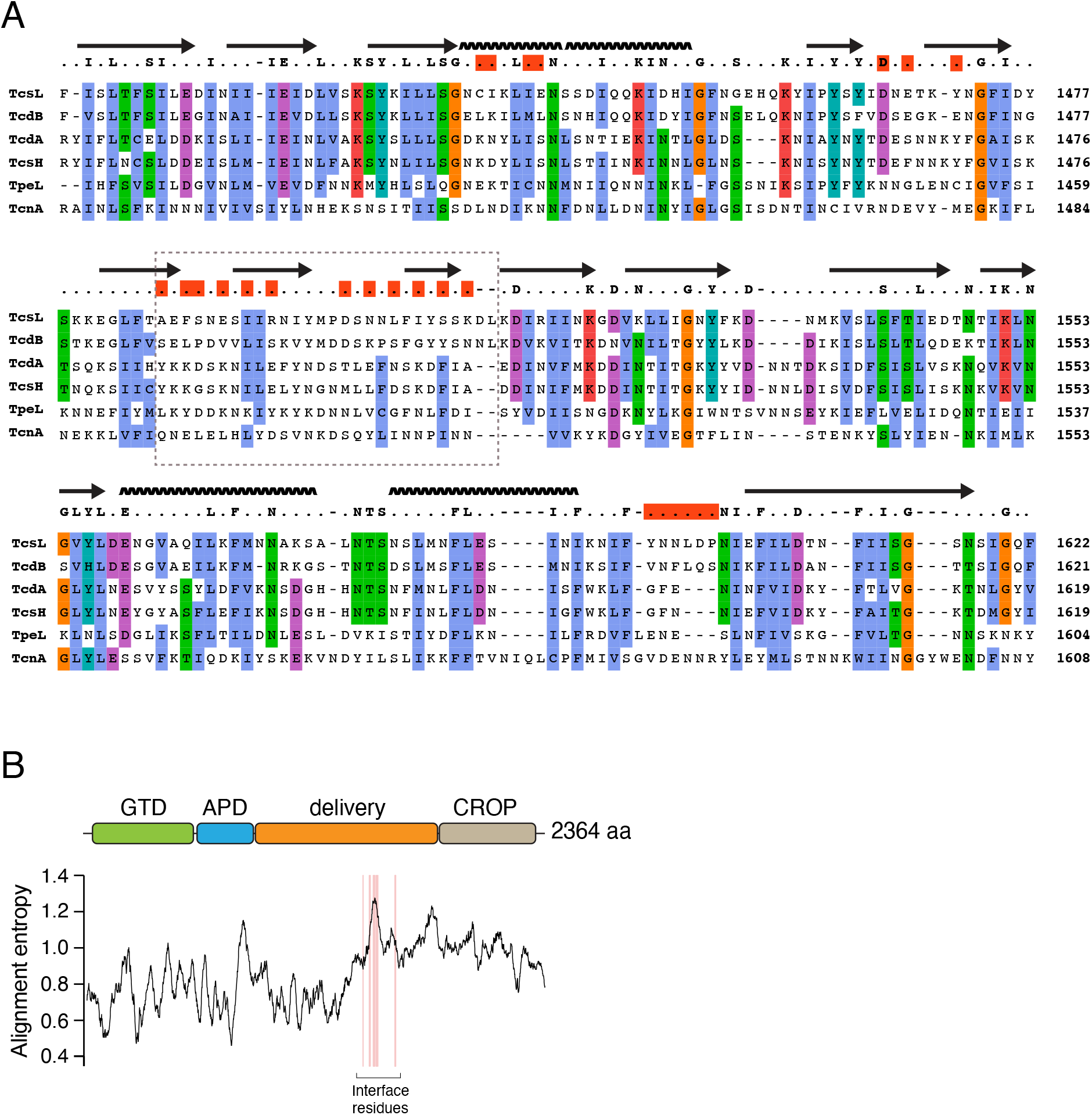
The receptor-binding surface of TcsL and TcdB is highly divergent between all clostridial toxins. (A) Partial sequence alignment of six known large clostridial toxins. Secondary structure elements and the consensus sequence (at least 4/6 identical residues) are shown above the alignment. Amino acids are colored based on their biophysical properties (ClustalX coloring) if at least 4/6 residues are similar in each column. Red boxes indicate interface residues in TcsL/SEMA6A and TcdB/Fzd7 complexes. The grey box highlights evolutionarily divergent beta sheets in the receptor-binding interface. (B) Sequence alignment entropy in large clostridial toxin family. The plot shows a moving average of alignment entropy (40-aa window around the central amino acid). Note that the highest entropy peak is in the receptor-binding interface. TcsL/SEMA6A and TcdB/FZD2 interface residues are indicated in light red. Top, domain organization of TcdB.

**Table S1.** MAGeCK scores from CRISPR/Cas9 screens in Hap1 cells with 1 nM and 0.1 nM TcsL.

**Table S2.** Cryo-EM data collection, image processing and model building statistics.

| **Data Collection** | TcsL-SEMA6A | SEMA6A |
| --- | --- | --- |
| Electron microscope | Titan Krios G3 |  |
| Camera | Falcon 4EC |  |
| Voltage (kV) | 300 |  |
| Nominal magnification | 75,000 |  |
| Calibrated physical pixel size (Å) | 1.06 |  |
| Total exposure (e- /Å<sup>2</sup>) | 45.24 |  |
| Number of frames | 30 |  |
| **Image Processing** |  |  |
| Motion correction software | Patch Motion/cryoSPARCv2 |  |
| CTF estimation software | Patch CTF/cryoSPARCv2 |  |
| Particle selection software | cryoSPARCv2 |  |
| 3D map classification and refinement software | cryoSPARCv2 |  |
| Micrographs used | 4581 |  |
| Particles selected | 3 896 638 |  |
| Global resolution (Å) | 3.3 | 3.1 |
| Particles contributing to final maps | 155,353 | 229,275 |
| **Model Building** |  |  |
| Modeling software | Coot, phenix.real_space_refine |  |
| Number of residues built | 1240 | 1042 |
| RMS (bonds) | 0.003 | 0.003 |
| RMS (angles) | 0.561 | 0.709 |
| Ramachandran favoured (%) | 93.4 | 94.1 |
| Rotamer outliers (%) | 0.64 | 0.00 |
| Clashscore | 3.58 | 2.33 |
| MolProbity score | 1.58 | 1.41 |

**Table S3.** TcsL residues contacting SEMA6A. BSA – buried surface area.

| Residue Number | Residue Type | BSA (Å<sup>2</sup>) |
| --- | --- | --- |
| 1433 | CYS | 6 |
| 1434 | ILE | 90 |
| 1435 | LYS | 2 |
| 1437 | ILE | 2 |
| 1438 | GLU | 23 |
| 1471 | TYR | 12 |
| 1484 | PHE | 1 |
| 1486 | ALA | 10 |
| 1488 | PHE | 63 |
| 1489 | SER | 9 |
| 1491 | GLU | 27 |
| 1493 | ILE | 43 |
| 1495 | ARG | 43 |
| 1505 | LEU | 15 |
| 1507 | ILE | 21 |
| 1509 | SER | 12 |
| 1511 | LYS | 11 |
| 1596 | TYR | 73 |
| 1597 | ASN | 51 |
| 1598 | ASN | 11 |
| 1599 | LEU | 18 |
| 1600 | ASP | 48 |
| 1601 | PRO | 31 |

**Table S4.** SEMA6A residues contacting TcsL.

| Residue Number | Residue Type | BSA (Å<sup>2</sup>) |
| --- | --- | --- |
| 102 | ALA | 29 |
| 105 | ASP | 38 |
| 106 | THR | 53 |
| 108 | ARG | 74 |
| 109 | MET | 141 |
| 110 | LYS | 107 |
| 111 | GLY | 26 |
| 141 | ASN | 10 |
| 189 | ASP | 19 |
| 190 | PHE | 33 |
| 191 | LEU | 75 |
| 193 | ILE | 28 |
| 212 | HIS | 32 |

**Movie S1. 3D variability analysis of TcsL.**

